# New tools reveal PCP-dependent polarized mechanics in the cortex and cytoplasm of single cells during convergent extension

**DOI:** 10.1101/2023.11.07.566066

**Authors:** Shinuo Weng, Caitlin C. Devitt, Bill M. Nyaoga, Anna E. Havnen, José Alvarado, John B. Wallingford

## Abstract

Understanding biomechanics of biological systems is crucial for unraveling complex processes like tissue morphogenesis. However, current methods for studying cellular mechanics *in vivo* are limited by the need for specialized equipment and often provide limited spatiotemporal resolution. Here we introduce two new techniques, Tension by Transverse Fluctuation (TFlux) and *in vivo* microrheology, that overcome these limitations. They both offer time-resolved, subcellular biomechanical analysis using only fluorescent reporters and widely available microscopes. Employing these two techniques, we have revealed a planar cell polarity (PCP)-dependent mechanical gradient both in the cell cortex and the cytoplasm of individual cells engaged in convergent extension. Importantly, the non-invasive nature of these methods holds great promise for its application for uncovering subcellular mechanical variations across a wide array of biological contexts.

**Summary:** Non-invasive imaging-based techniques providing time-resolved biomechanical analysis at subcellular scales in developing vertebrate embryos.

## Introduction

Individual cells both generate and sense mechanical forces and the crucial role of single-cell mechanics in the larger-scale emergent processes of tissue morphogenesis is now well established. That said, the general principles of individual cell mechanics *in vivo* remain poorly defined, and several hurdles stand between us and a more comprehensive understanding. First, we currently possess only a sparse sampling of subcellular mechanics in embryos, with most work focused on a relatively narrow set of cell types and organisms. This limitation stems in part from the sophisticated hardware required for existing methods. For example, while atomic force microscopy (AFM), pulsed laser systems, optical trapping, micro-aspiration, and magnetic manipulation of ferrofluids are the core tools of embryo biomechanics analysis *in vivo* (Campàs, 2016; Davidson, 2011; Gjorevski and Nelson, 2010; Haase and Pelling, 2015; Trier and Davidson, 2011; Trubuil et al., 2021), relatively few investigators have access to such instrumentation.

Alternative, imaged-based methods collectively termed “force inference” have been developed (Roca-Cusachs et al., 2017; Roffay et al., 2021; Veldhuis et al., 2015), but these generally rely on cell geometry and force balance across multiple cells in any given image at one time point, which introduce two key limitations: The methods have difficulty assessing *dynamic* changes in cell cortex tension over time and they generally do not provide information on *sub-cellular* heterogeneity of cortex tension.

This second issue is also a more general hurdle, as many commonly used methods do not provide sufficient spatial resolution to address the issue of sub-cellular mechanics. For example, the widely used method of laser microdissection generally reports only the mean behavior of the entire junction. In the realm of theory, most commonly used vertex models of morphogenesis also treat junctions as mechanically homogeneous along their lengths (Alt et al., 2017; Fletcher et al., 2014). These are key concerns because evidence from cultured cells, from embryos, and from theory all suggest that individual cell-cell junctions can display significant mechanical heterogeneity (Cavanaugh et al., 2022; Huebner et al., 2021b; Lieber et al., 2015; Shi et al., 2018; Strale et al., 2015; Vanderleest et al., 2018).

Finally, while we have witnessed an acceleration of recent work on cell cortex mechanics in embryonic morphogenesis (e.g. (Heer and Martin, 2017; Lecuit and Lenne, 2007; Stooke-Vaughan and Campàs, 2018), the mechanical properties of the cytoplasm also influence cell behavior (Mathieu and Manneville, 2019; Salbreux et al., 2012; Wirtz, 2009). For example, local patterning of cytoplasmic stiffness is important for the movement of fibroblasts and axonal growth cones *in vitro* (Kole et al., 2005; O′Toole et al., 2008). *In vivo*, hydraulic behavior of the cytoplasm transmits forces generated at cell-cell junctions to form the *Drosophila* ventral furrow (He et al., 2014), and cytoplasmic pressure in *Drosophila* germ cells tunes the shape of neighboring follicle cells (Lamiré et al., 2020). Nonetheless, apart from initial samplings in the early embryos of *Drosophila* and *C. elegans* (Daniels et al., 2006; Doubrovinski et al., 2017), very few studies of cytoplasmic viscosity in embryonic morphogenesis have been reported. Moreover, the relationship between the cytoplasmic and cortical mechanical properties remains an uncharted territory. New tools for the exploration of these important issues would therefore be welcome.

Here, we report two new approaches for quantifying the mechanical properties of the cell cortex and the cytoplasm in vertebrate embryonic cells. Each method enables *1)* time-resolved biomechanics *in vivo, 2)* at subcellular scales, *3)* using only fluorescent reporters and widely available microscopes. Using these tools, we describe and quantify planar polarized mechanics of both the cortex and the cytoplasm in individual cells engaged in convergent extension during *Xenopus* gastrulation and show that both are dependent on planar cell polarity (PCP) signaling. Critically, because these tools require only expression of fluorescent proteins and reasonably high spatiotemporal imaging resolution, they should be easily adaptable to other systems, and thus have the potential to provide a far broader view of the diversity of cellular and subcellular biomechanics during embryonic morphogenesis.

## Results

Our desire to develop new methods for single cell biomechanics *in vivo* was spurred by studies of cell intercalation during convergent extension (CE), the evolutionarily conserved collective cell movement that elongates diverse tissues and organs (Huebner and Wallingford, 2018; Tada and Heisenberg, 2012). In a previous study, we showed that the transverse fluctuations of vertices provide a convenient proxy for local mechanics (Huebner et al., 2021b). We therefore reasoned that quantifying transverse fluctuation at *all pixels* along a junction might provide a temporally and spatially resolved reporter of mechanical heterogeneity (**Fig. 1A)**.

**Figure 1.**
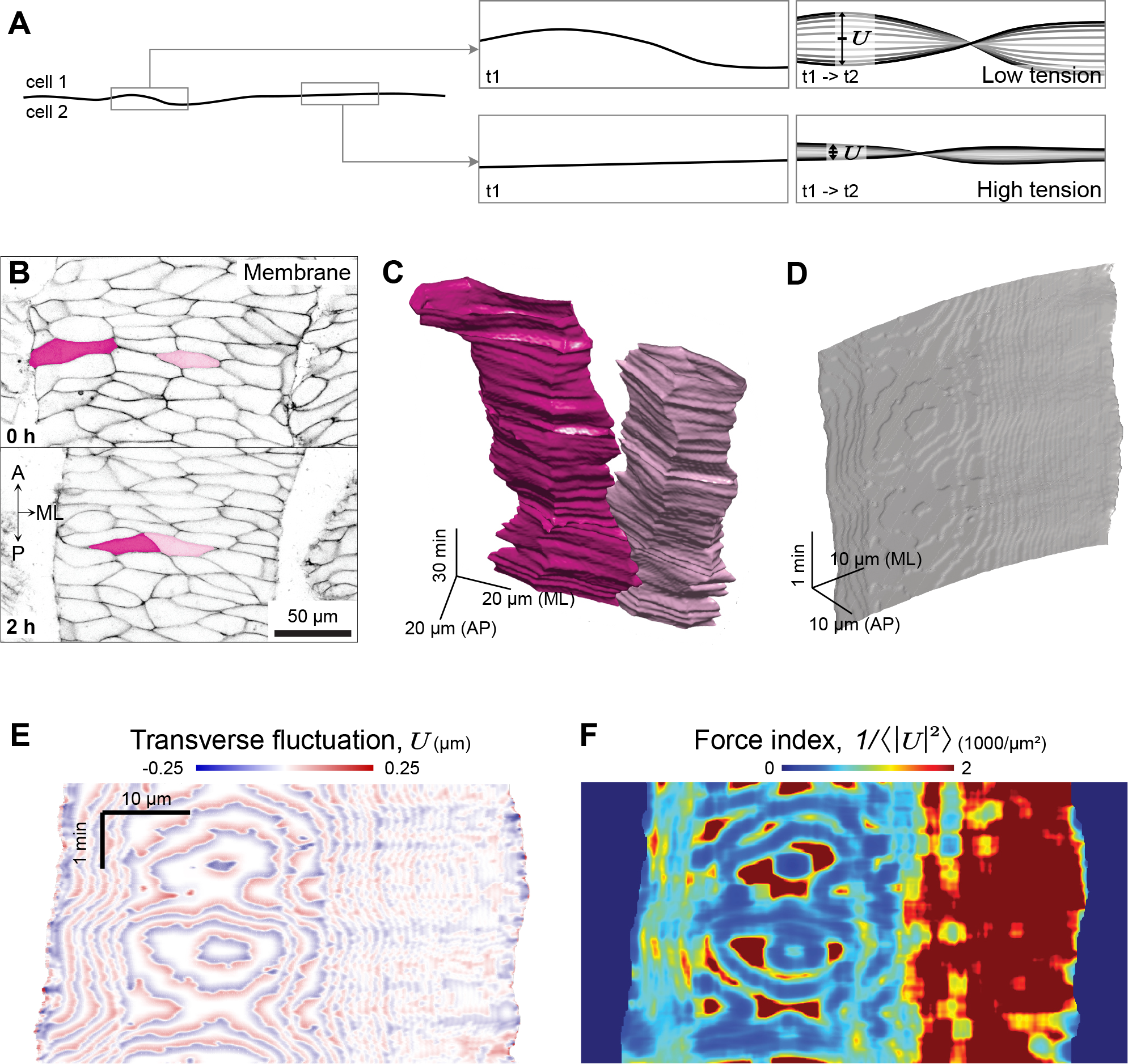
The TFlux Concept and Method. **A**. Schematic of TFlux concept. **B**. Still frames from a time- lapse movie with two cells indicated. **C**. 3D (X-Y-Time) view of segmented volumes of the two cells indicated in panel B. **D**. 3D (X-Y-Time) view of a single segmented junction. **E**. Kymographic heatmap of the junction shown in D with time and mediolateral position indicated in the kymograph and transverse fluctuation indicated by the heat map. **F**. Kymographic heatmap as shown in E, but with the heatmap indicating force index.

### TFlux allows subcellular, time-resolved assessment of cell cortex tension *in vivo*

To visualize transverse fluctuations of cell cortices, we labeled membranes with a fluorescent protein tagged with CAAX, and we imaged cells in the *Xenopus* gastrula mesoderm with confocal microscopy. We then segmented cells with Ilastik^®^ pixel classification and carving (Berg et al., 2019; Kreshuk and Zhang, 2019) to generate 3D (X-Y-time) meshes (**Fig. 1B, C**). By taking the microscopy at 0.31 μm/pixel and 0.05 frame/sec, the coarse meshes revealed highly heterogeneous cell behaviors associated with overall cell movement (**Fig. 1C**). We then increased both spatial and temporal resolution to 0.205 μm/pixel and 1.0 frame/sec for microscopy, followed the same procedure to generate finer 3D meshes, and used a customized MATLAB script to focus our analysis on junction of interest over time (**Fig. 1D**).

To quantify transverse fluctuations, we defined a base-line position for each junction at each time point by calculating a moving average in space over 2 μm along the length of the junction and in time over 20 seconds (**Supp. Fig. 1A-C**). The selection of these specific baseline sampling rates was guided by two key considerations: *1)* In the spatial domain, we aimed to capture local mechanical heterogeneity imparted by cadherin clusters, which typically act at the 1 μm scale (Huebner et al., 2021a). Hence, we opted for a 2 μm distance for background smoothing. *2)* In the temporal domain, we wanted to eliminate overall cell movement as shown in **Fig. 1C**, so we employed the same 20-sec interval for temporal background smoothing. Transverse fluctuation (TFlux) was then measured at each time point as the distance from the junction position to the baseline (**Supp. Fig. 1D**).

Consistent with our previous findings with multicellular vertices (Huebner et al., 2021b), Kymographic heatmaps demonstrate that cell-cell junctions displayed localized, heterogeneous patterns of transverse fluctuation (**Fig. 1E**). From these data, we defined a “force index” as the reciprocal of the mean square transverse fluctuation (Alvarado and Koenderink, 2014; MacKintosh et al., 1995) of the junction over 2 μm along the length and over a 20-second period of observation (**Fig. 1F**). This in turn allows us to quantify cell cortex tension in both space and time (**Fig. 1F**). Importantly, this technique reproduced the mediolateral mechanical heterogeneity reported in our previously work (Huebner et al., 2021b), but now with additional spatiotemporal insights that warrant further exploration.

To validate TFlux as a relevant metric, we reasoned that cell cortex tension is imparted largely by a contractile actomyosin network just below the plasma membrane, and that disruption of actomyosin should reduce the TFlux force index. We therefore pharmacologically inhibited myosin using Blebbistatin or actin using Latrunculin B (Morton et al., 2000; Straight et al., 2003; Wakatsuki et al., 2001) and made paired comparisons of the same junction before and aSer addition of drug (**Fig. 2A, a’**). To also test the sensitivity of TFlux, we used doses of these drugs well below those required to robustly disrupt CE (Devitt et al., 2023; Skoglund et al., 2008). Critically, even though these low doses elicited no apparent changes in cell shape (**Fig. 2B, b’**), they did significantly reduce the force index of junctions (**Fig. 2C, D**).

**Figure 2.**
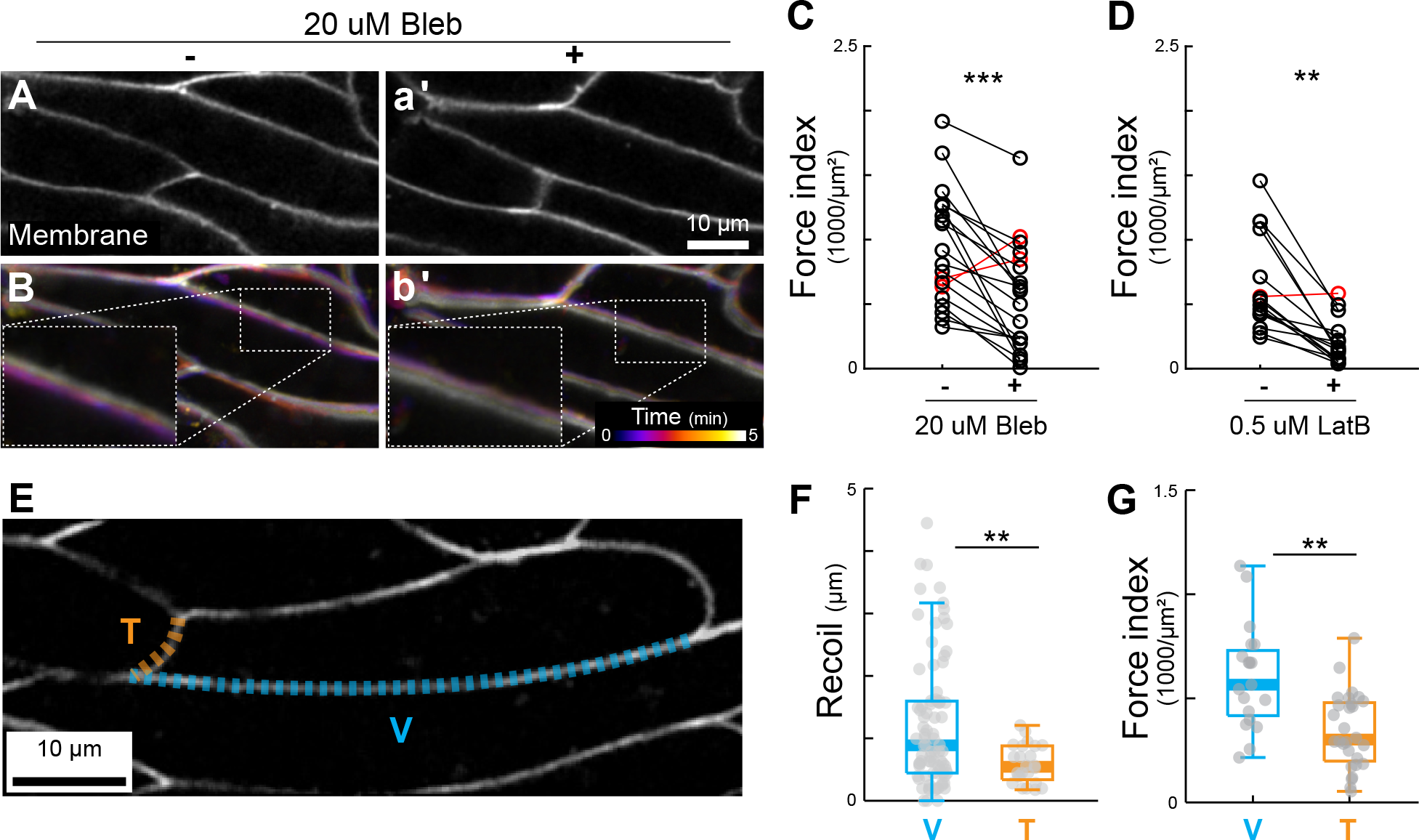
Valida1on of TFlux Method. **A, a’**. Still images from time lapse of cells engaged in CE before and aSer addition of blebbistatin. **B, b’**. Higher magnification images of a single junction, color-coded for time to reflect overall junction movement, before and aSer addition of blebbistatin. **C**. Paired TFlux measurements of single junctions before and aSer addition of blebbistatin. **D**. Paired TFlux measurements of single junctions before and aSer addition of Latrunculin B. **E**. Still image from time lapse of CE with a mediolaterally aligned “v-junction” and an anteroposteriorly aligned “t-junction” indicated. **F**. Laser microdissection data from Shindo and Wallingford, 2014 showing elevated cortex tension in v-junctions. **G**. TFlux data showing elevated cortex tension in v-junctions.

As a further test, we asked if TFlux could recapitulate known differences in cell cortex tension during CE. We previously showed using laser microdissection that mediolaterally-aligned “v-junctions” display roughly 2.1-fold higher cortical tension than do anteroposteriorly aligned “t-junctions” (**Fig. 2E, F**)(Shindo and Wallingford, 2014). Using TFlux, we observed a similar (1.7-fold) difference in these tensions (**Fig. 2G**). Together, the ability to detect both the effects of actomyosin disruption and known tension differences determined by laser microdissection suggest that TFlux is a sensitive and useful method for *in vivo* analysis of cell cortex mechanics.

### TFlux reveals a PCP-dependent polarity in cell cortex tension within single cells enaged in convergent extension

To ask if TFlux could provide new insights, we considered that CE requires a superficial “node-and-cable” meshwork of actin (Kim and Davidson, 2011; Pfister et al., 2016; Skoglund et al., 2008). Because we recently found this meshwork to be organized differently in the anterior and posterior regions of single cells (Devitt et al., 2023), we used TFlux to test the possibility that the anterior and posterior cell cortices of each individual cell during CE may experience distinct tensions.

We labeled the plasma membrane of single cells in an otherwise unlabeled notochord using targeted microinjection (Kieserman et al., 2010), segmented the anterior and posterior cell membrane, and performed TFlux (**Fig. 3A**). By taking paired measurements from multiple individual cells, we found that the anterior plasma membrane consistently displayed significantly elevated cortical tension (i.e., less transverse fluctuation and high force index) compared to the posterior of the same cell (**Fig. 3B, C**).

**Figure 3.**
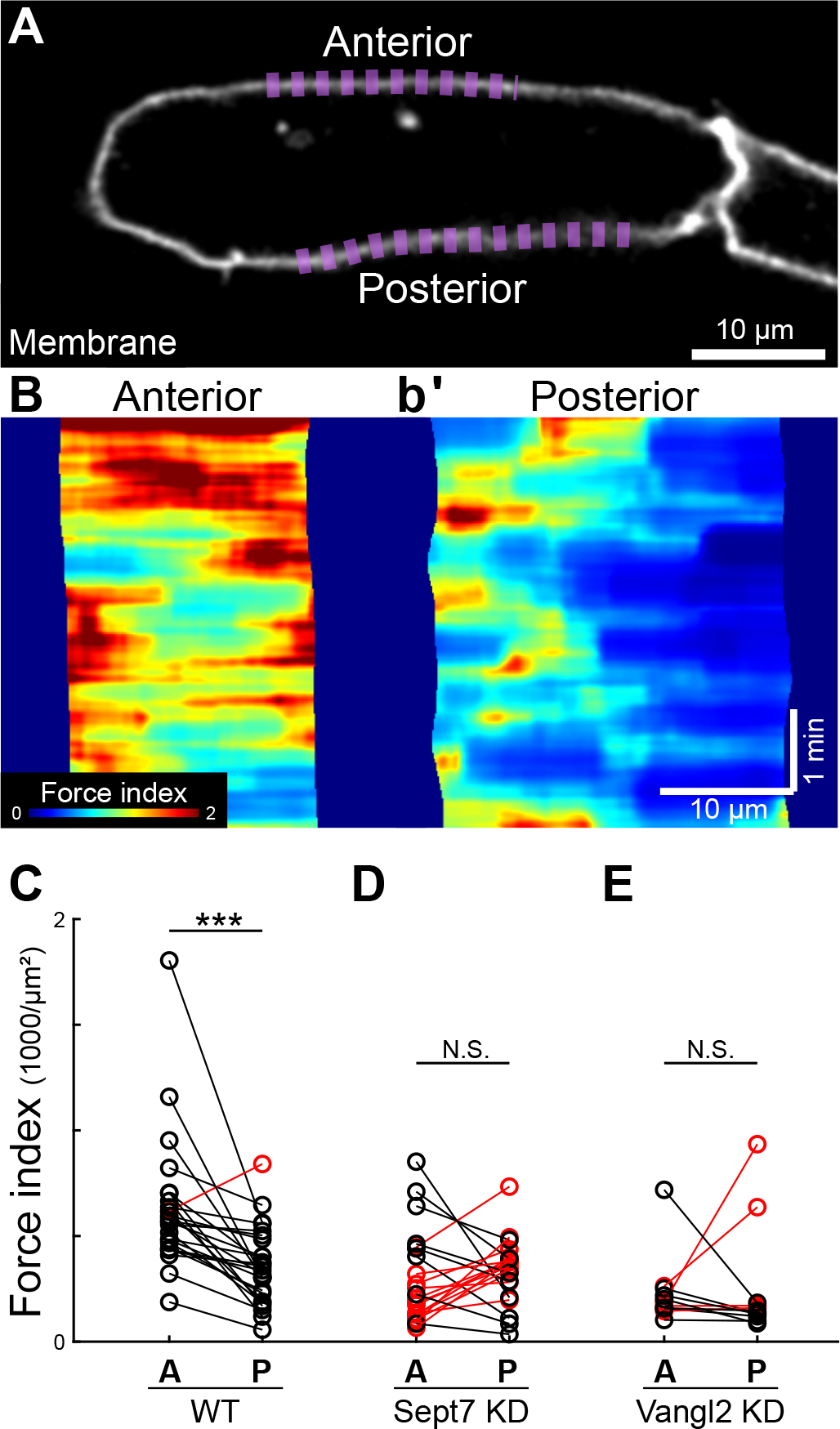
Anterior and posterior faces of cells display PCP-dependent differences in cell cortex tension during CE. **A**. Still image of a single labelled cells in a background of unlabeled cells, all engaged in CE, with anterior and posterior cell faces indicated. **B, b’**. Kymographic heatmap indicating force index for the anterior and posterior cell faces indicated in panel A. **C-E**. Quantification of mean force index for anterior and posterior cell faces for control cells, Sept7-KD cells, and Vangl2-KD cells.

In our previous study, we also found that the anteroposterior polarization of actin node-and-cable organization requires the core PCP protein Vangl2 and its downstream effector, the actin regulator Septin7 (Devitt et al., 2023). Consistent with a link between the polarity of actin organization and the polarity of cell cortex tension, we found that knockdown of either Vangl2 or Septin7 eliminated the differences in cell cortex tension between the anterior and posterior faces of single cells (**Fig. 3D, E**). Interestingly, loss of Vangl2 or Septin7 disrupted polarization by reducing the elevated tension anteriorly (i.e., by causing anterior tensions in experimental cells to resemble posterior tensions in control cells)(**Fig. 3D, E**). It’s noteworthy then that Vangl2 or Sept7 KD had a parallel effect on actin organization (Devitt et al., 2023), correlating higher order of cytoplasmic actin organization with higher cortical tension.

### A phase separated fluorescent report enables *in vivo* microrheology

Combined with an accelerating appreciation of local mechanical heterogeneities in cell cortex mechanics (Cavanaugh et al., 2022; Huebner et al., 2021b; Vanderleest et al., 2018), our findings with TFlux prompted us to ask if cells displayed similar subcellular heterogeneity in the mechanics of the cytoplasm. We therefore turned to microrheology, which is typically performed with fluorescent microbeads that are tracked over time providing measurement of mean-squared displacement (MSD) that in turn reports cytoplasmic viscosity (Weihs et al., 2006; Wirtz, 2009). The method has been applied in several cultured cell types (Daniels et al., 2006; Kole et al., 2005; Weihs et al., 2006; Wirtz, 2009), as well as in *Drosophila* and *C. elegans* embryos (Daniels et al., 2006; He et al., 2014). However, in the latter cases, analysis was limited to only early stages and few results have been reported. In *Xenopus*, we found that microbeads clump and are extruded (not shown).

We reasoned that phase-separated, droplet-forming proteins may provide a solution (Banani et al., 2017; Shin and Brangwynne, 2017). Indeed, expression of a fluorescent fusion to the phase-separating stress granule protein Tia1 (Mackenzie et al., 2017) generated discrete, well-dispersed foci in the cytoplasm *Xenopus* notochord cells (**Fig. 4A**). Foci remained brightly fluorescent when imaged at 2 frames per second over approximately 2 minutes, allowing effective tracking (**Fig. 4B, b’**).

**Figure 4.**
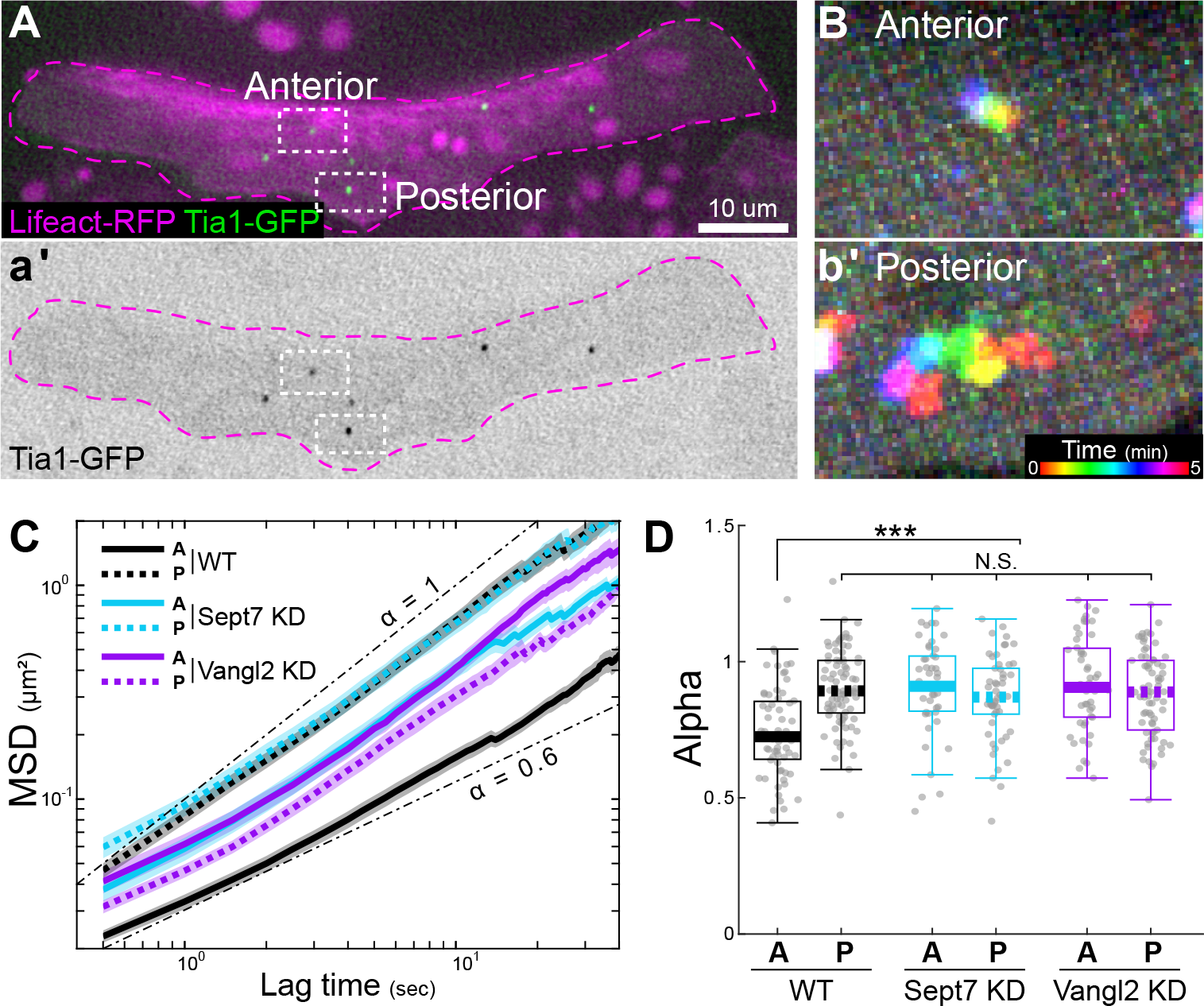
Cytoplasmic microrheology with a phase-separated reporter reveals PCP-dependent differences in cytoplasmic mechanics during CE. **A**. Still image of a single cell expressing Tia1-GFP and Lifeact-RFP. **a’**. Tia1GFP-only channel. **B**. Time-coded image showing minimal movement of a single Tia1-GFP particle in the anterior cytoplasm. **b’**. Time-coded image showing substantial movement of a single Tia1-GFP particle in the posterior cytoplasm. **C**. Mean Squared Displacement for anterior (solid) and posterior (dashed) particles in control (black), Sept7-KD (blue) and Vangl2-KD (magenta) cells. **D**. Mean anomalous exponent α for the MSD of particles in control, Sept7-KD (blue), and Vangl2-KD (magenta) cells.

Using time-lapse movies of phase separated foci, we measured MSDs as a function of lag time (**Fig. 4C**). We performed a rough estimation of the cytoplasmic viscosity, by assuming the Brownian motion of the Tiati-GFP foci, where the MSD of a random walker scales linearly with time, i.e. diffusion (Weihs et al., 2006; Wirtz, 2009). We estimated a mean cytoplasmic viscosity of 2.20 ± 0.20 Poise. Importantly, this values is generally consistent with viscosities observed for the cytoplasm of one-cell *C. elegans* embryos or serum starved 3T3 fibroblasts (Daniels et al., 2006; Kole et al., 2005; Wirtz, 2009), suggesting our method is generally useful for assessing cytoplasmic viscosity *in vivo*.

### A PCP-dependent gradient of cytoplasmic viscosity in single cells enaged in convergent extension

We also noticed that the diffusive motion of Tia1-GFP foci was anomalous, in that the MSD follows a sublinear increase with time, fiwng a power-law relation with the exponent α ti 1. Such a subdiffusive motion has been proposed as a measure of macromolecular crowding in the cytoplasm (Weiss et al., 2004), and recent work has directly linked cytoplasmic crowding with sub-diffusivity in both yeast and human cells (Delarue et al., 2018).

We therefore examined the anomalous diffusion exponent α to ask if the macromolecular crowding of the cytoplasm of cells engaged in CE displayed a polarization similar to that observed for the cell cortex (**Fig. 3**) and the actin cytoskeleton (Devitt et al., 2023). We found that Tia1-GFP-labelled foci in the anterior regions of cells consistently displayed highly constrained sub-diffusion, with an exponent α ∼ 0.7(**Fig. 4C, D**). By contrast, particles in the posterior region displayed a significantly more diffusive movement, with the exponent α closer to 1 (**Fig. 4C, D**). These data suggest that the anterior cytoplasm of a cell engaged in CE is more crowded than the posterior cytoplasm of the same cell.

Strikingly, knockdown of either Vangl2 or Sept7 eliminated these anterior-to-posterior differences in particle motion, suggesting a compromised anterior-to-posterior polarity in the internal organization of the cell (**Fig. 4C, D**). These effects were not related to any change in the size of Tia-GFP foci in the different conditions (**Supp. Fig. 2**). Vangl2 or Sept7 KD specifically reduced the normally elevated viscosity in anterior regions, reflecting similar effect on cortex tension (**Fig. 3**) and actin organization (Devitt et al., 2023) in the anterior regions of these cells. Thus, disruption of PCP signaling or its downstream effector Sept7 results in loss of an anteroposteriorly oriented gradient of actin organization, cell cortex tension, and cytoplasmic viscosity in individual cells engaged in CE.

## Discussion

Here, we describe new tools for assessing sub-cellular biomechanics *in vivo*. We deploy these tools to study convergent extension, a process that was the subject of many seminal studies of tissue mechanics (Adams et al., 1990; Feroze et al., 2015; Jacobson and Gordon, 1976; Moore et al., 1995; Shook et al., 2018; von Dassow and Davidson, 2009; Zhou et al., 2009; Zhou et al., 2010). More recently, the underlying mechanics of single cells involved in CE have begun to be probed (Huebner et al., 2021b; Shawky et al., 2018; Shindo and Wallingford, 2014), but those studies have been limited, in part due to the limitations of existing methods. Using TFlux and *in vivo* microrheology, we now provide evidence for a PCP-dependent mechanical gradient across each single cell in the *Xenopus* notochord during CE. The mechanical gradient was observed using tools to probe both the cytoplasm and the cell cortex.

These data are significant for several reasons. First, they provide a link between the molecular and the mechanical properties at a subcellular level during vertebrate CE. Indeed, the core PCP proteins are enriched on the anterior and posterior faces of cells during CE (Butler and Wallingford, 2018; Ciruna et al., 2006; Shindo et al., 2019; Yin et al., 2008). These, in turn, impart an anteroposterior organization of the actin cytoskeleton (Devitt et al., 2023), and our data here suggest that this in turn imparts a mechanical gradient. Second, this coherent molecular/mechanical gradient adds weight to the idea that very local, sub-cellular mechanical heterogeneities are important features of cell movement, both in single cultured cells and in tissues (Cavanaugh et al., 2022; Huebner et al., 2021b; Lieber et al., 2015; Shi et al., 2018; Strale et al., 2015; Vanderleest et al., 2018). Third, our data suggested that cells are mechanically polarized in two axes, both mediolaterally and anteroposteriorly. The functions of such a cellular mechanical pattern warrant further exploration.

Finally, the methods reported here offer broad utility. Their non-invasive nature allows the collection of time-resolved data, making them especially attractive for studies of cell movement. Moreover, these techniques require only expression of fluorescent proteins and microscopes with reasonable temporal and spatial resolution, and so should be easily adaptable by many labs to many biological systems. Moreover, the methods are also compatible with other image-based approaches, such as assessment of cadherin dynamics using fluorescence recovery aSer photobleaching (FRAP)(Huebner and Wallingford, 2022) or membrane tension through fluorescence lifetime imaging microscopy (FLIM)(Colom et al., 2018).

Of particular interest, the TFlux method opens doors to a more comprehensive understanding of biomechanical and biophysical features in tissue morphogenesis. In our current analysis, we have chosen specific baseline smoothing rates to quantify transverse fluctuation. The potential for a comprehensive spatial and temporal analysis of the 3D mesh in **Fig. 1C** holds great promise for identifying additional matrices linked to various cellular and subcellular behaviors and properties. Furthermore, while our force index is presently defined based on a model of a semiflexible biopolymer at equilibrium pulled at two ends, it is important to acknowledge that cell cortex experiences forces along its entire length with variations. The method’s utility could be further enhanced with a more precise biophysical model. Our aspiration is that by extending these analyses across multiple scales, and exploring diverse tissue and animal types, we come closer to a more comprehensive understanding of biomechanics in tissue morphogenesis.

## Acknowledgements

This work was supported by R01HD099191 and R21HD112657.

## Figure Legends

## Supplemental Materials and Methods

### *Xenopus* embryo manipula1ons

Ovulation was induced by injecting adult female *Xenopus* laevis with 600 units of human chorionic gonadotropin (HCG, MERCK Animal Health) and animals were kept at 16 dc overnight. Eggs were collected the following day by gently squeezing the ovulating females, and were fertilized *in vitro*. Eggs were dejellied in 2.5% cysteine (pH 7.9) 1.5 hours aSer fertilization and moved to 1/3x Marc’s modified Ringer’s (MMR) media. For micro-injection, embryos were placed in 2% ficoll in 1/3x MMR or DI Water during injection and moved to 1/3x MMR 30 min aSer injection. Embryos were injected in the dorsal blastomeres at the 4-cell stage targeting the C1 cells at 32-cell stage and presumptive notochord. For mosaic labelling, embryos were injected at the 32-cell stage targeting one of the C1 cells. Keller explants were dissected at stage 10.25 in Steinberg’s solution using hair tools.

### Plasmids and mRNA for microinjec1ons

Membrane-BFP, membrane-GFP, membrane-RFP plasmids were made in pCS2+, Lifeact-RFP was made in pCS, and Tia1-GFP was made in pCS10R. Capped mRNAs were generated using the ThermoFisher SP6 mMessage mMachine kit (Catalog number: AM1340). The amount of injected mRNAs per blastomere were as follows: Membrane-BFP, membrane-GFP, membrane-RFP or Lifeact-RFP, 60-80 pg per injection at 4-cell stage or 8-10 pg per injection at 32-cell stage; Tia1-GFP, 50 ng per injection at 4-cell stage or 6 ng per injection at 32-cell stage.

### Inhibitor treatments

Inhibitors were solubilized in DMSO to a stock concentration of 2 mM for Latrunculin B (Cayman Chemical, 10010631) and 10 mM for para-amino-Blebbistatin (Cayman Chemical, 22699). Explants were treated with 0.5 μM Latrunculin B or 20 μM of Blebbistatin diluted in Steinberg’s solution. Same samples were imaged before and 30 min aSer the addition of inhibitors.

### Imaging *Xenopus* explants

Explants were submerged in Steinberg’s solutions and cultured on glass coverslips coated with Fibronectin (Sigma-Aldrich, F1141) at 2 μg/cm^2^. All images for TFlux were taken on a laser scanning confocal microscope (Nikon A1R), at the focal plane 5 μm deep into the explant, and at the spatial and temporal resolution of 0.205 μm/pixel and 1.0 frame/sec, respectively. Images for microrheology were taken on a spinning disk confocal microscope (Nikon W1), at the spatial and temporal resolution of 0.130 μm/pixel and 2 frame/sec, respectively.

### Tension by Transverse Fluctua1on (TFlux)

TFlux measurement is based on the idea that the transverse fluctuation of a junction is inversely proportional to tension on the junction. To track the junction movement, we labeled cell membrane with fluorescent protein tagged with CAAX and imaged on a Nikon A1R confocal microscope for 5 min at the spatial and temporal resolution of 0.205 μm/pixel and 1.0 frame/sec, respectively. Prior to segmentation, we preprocessed the timelapse movies in ImageJ following the flowing procedures to improve the cost efficiency of segmentation: crop region of interest, smooth once, subject background, and adjust contrast without oversaturation. ASer that, preprocessed time-lapse movies were imported into Ilastik® and cells of interest were segmented using Ilastik® pixel classification and Ilastik® Carving. 3D (xyt) meshes of cells were then imported to a customized MATLAB script to detect the transverse fluctuation of regions of interest over time. Briefly, we first defined a base-line position of the junction at each time point to account for the overall movement of the cells and the junction. To do so, we performed a moving average over 2 μm along the junction length at each time point (**Supp. Fig. 1B**), and then another moving average over a 20-sec period in time (**Supp. Fig. 1C**). Transverse fluctuation was then measured at each time point as the distance from the original junction position to the base line (**Supp. Fig. 1D**). We defined a “force index” as the reciprocal of the mean square transverse fluctuation (Alvarado and Koenderink, 2014) of the junction. For the kymographic heatmap, force index was calculated over 2 μm along the length of the junction and over a 20-second period of observation. For the overall junction tension, force index was calculated along the entire junction over a 5-min of observation.

### Particle tracking microrheology

Particle tracking microrheology was performed by tracking the displacement of Tia1-GFP particles using Nikon W1 spinning disk confocal microscope. Time-lapse movies of Tia1-GFP particles in cells co-expressing Lifeact-RFP were taken at the interval of 0.5 sec for 5 minutes. Customized MATLAB scripts were used to track the particle displacement, and then generate the Mean Square Displacement (MSD), ⟨r^2 (τ)⟩ as a function of lag time, τ for each particle. We presumed that the MSD data follows the anomalous diffusion and thus the MSD follows a power law ⟨r^2 (τ)⟩ ∝ τ^α. The power law exponent α=1 for Brownian motion; α ti1 for subdiffusion, a sign of macromolecular crowding; and α>1 for superdiffusion, when particle movement is strongly influenced by other active cellular movement, such as actin flow.

## Supplemental Figure Legends

**supp. Figure 1.**
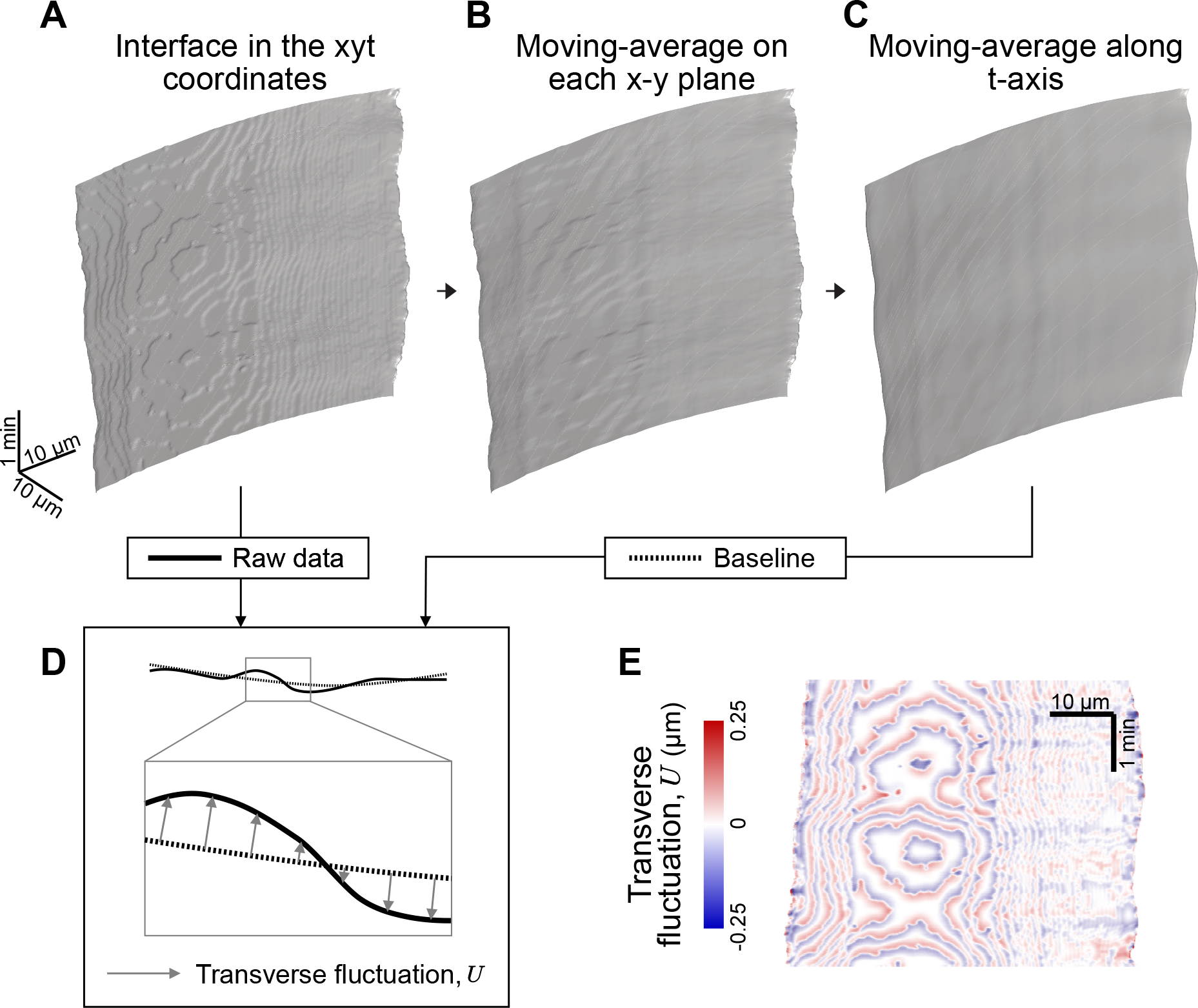
Measurement of transverse fluctua1on. **A**. 3D (X-Y-Time) view of a single segmented junction. **B**. 3D (X-Y-Time) view of the same junction as in A aSer a moving average over 2 μm along the junction length at each time point. **C**. 3D (X-Y-Time) view of the same junction as in B followed by a moving average over a 20-sec period. **D**. Transverse fluctuation defined as the distance from the original junction position in A to the base line position in C. **E**. Kymographic heatmap of transverse fluctuation.

**supp. Figure 2.**
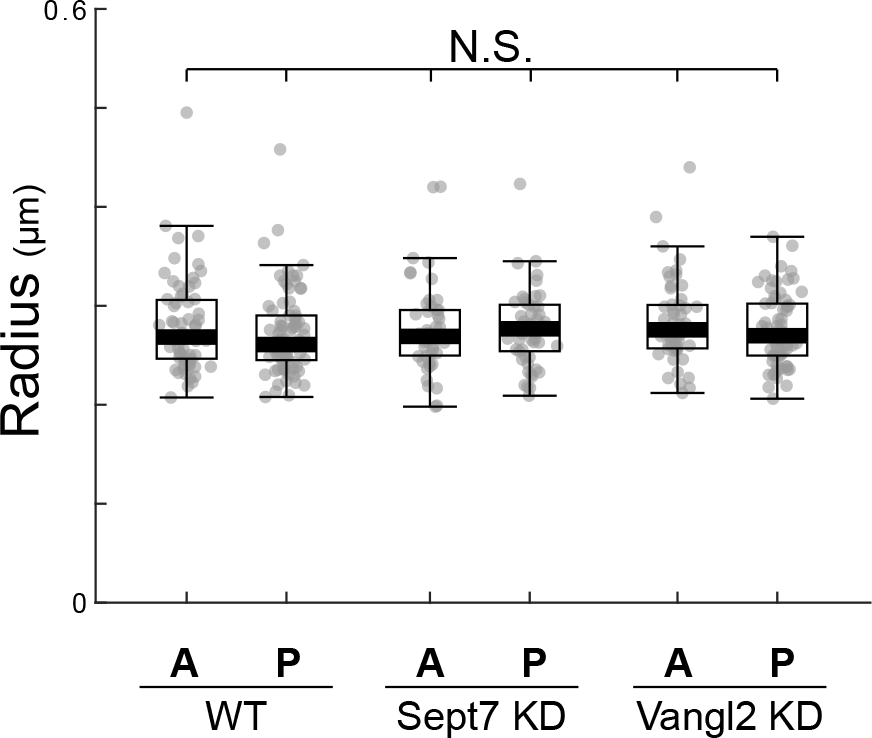
Size of Tia1 foci in different condi1ons.

